# Direct regulation of cell cycle regulatory gene expression by NtrX to promote *Sinorhizobium meliloti* cell division

**DOI:** 10.1101/2021.11.15.468759

**Authors:** Shenghui Xing, Lanya Zhang, Fang An, Leqi Huang, Xinwei Yang, Shuang Zeng, Ningning Li, Wenjia Zheng, Khadidja Ouenzar, Liangliang Yu, Li Luo

## Abstract

Cell division of the alfalfa symbiont, *Sinorhizobium meliloti*, is regulated by the CtrA signaling network. The gene expression of regulatory proteins in the network is affected by nutrient signaling. In this study, we found that NtrX, one of the regulators of nitrogen metabolic response, can directly regulate the expression of several regulatory genes from the CtrA signaling network. Three sets of *S. meliloti ntrX* mutants, including the plasmid insertion strain, the depletion strain and the substitution of the 53^rd^ aspartate (*ntrX^D53E^*) from a plasmid in the wild-type strain (Sm1021), showed similar cell division defects, such as slow growth, abnormal morphology of partial cells and delayed DNA synthesis. Transcript quantitative evaluation indicated that the transcription of genes such as *ctrA* and *gcrA* was up-regulated, while the transcription of genes such as *dnaA* and *ftsZl* was down-regulated in the insertion mutant and the strain of Sm1021 expressing *ntrX^D53E^*. Correspondingly, inducible transcription of *ntrX* activates the expression of *dnaA* and *ftsZ1*, but represses *ctrA* and *gcrA* in the depletion strain. The expression levels of CtrA and GcrA were confirmed by western blotting, which were consistent with the transcription data. The transcriptional regulation of these genes requires phosphorylation of the conserved 53^rd^ aspartate in the NtrX protein. The NtrX protein binds directly to the promoter regions of *ctrA*, *gcrA*, *dnaA* and *ftsZ1* by recognizing the characteristic sequence CAAN_2-5_TTG Our findings reveal that NtrX is a novel transcriptional regulator of the CtrA signaling pathway genes, and positively affects bacterial cell division, associated with nitrogen metabolism.

**IMPORTANCE:** *Sinorhizobium meliloti* infects the host alfalfa and induces formation of nitrogen-fixing nodules. Proliferation of rhizobia in plant tissues and cells is strictly controlled in the early stage of symbiotic interactions. However, the control mechanism is not very clear. Cell division of *S. meliloti* in the free-living state is regulated by the CtrA signaling network, but molecular mechanisms by which the CtrA system is associated with environmental nutrient signals (e.g., ammonia nitrogen) need to be further explored. This study demonstrates that NtrX, a regulator of nitrogen metabolism, required for symbiotic nodulation and nitrogen fixation by *S. meliloti* 1021, can act as a transcriptional regulator of the CtrA signaling system. It may link nitrogen signaling to cell cycle regulation in *Rhizobium* species.

## INTRODUCTION

*Caulobacter crescentus* is a model strain of α-proteobacteria in molecular cell biology (1). It takes advantage of one cell division to produce two cells with different shapes and sizes (2). In recent years, a complex cell cycle regulatory network has been revealed in this species. This network consists of multiple histidine kinases such as CckA, DivL, DivJ, and PleC, a histidine phosphotransfer protein ChpT, response regulators DivK and CpdR, and transcription regulators CtrA, GcrA, DnaA, SciP, and MucR (1–6). Among these cell cycle regulators, CtrA and GcrA negatively regulate cell division, which is opposite to DnaA. Although this network has been reported to possibly mediate nutritional signals for regulating bacterial growth and proliferation, the exact molecular mechanism is currently unclear.

*Sinorhizobium meliloti* is a model strain of rhizobia that can infect the host plant and form nitrogen-fixing nodules. During symbiosis, the cell division of *S. meliloti* on the surface of host plant alfalfa roots, at the front ends of extended infection threads and in the infection zones of root nodules, is stringently controlled (7), but how cell division is regulated is not very clear. Although NCR (Nodule Cysteine Rich) peptides secreted by host plants are known to induce terminal differentiation of bacteroids in host plant cells (8–11), several legumes do not produce NCR peptides. Therefore, there may be other mechanisms that control cell division of symbiotic rhizobia in host plants. Since *S. meliloti* and *C. crescentus* belong to α-proteobacteria, based on the research results of *C. crescentus*, with the aid of DNA sequence homology analysis, many cell cycle regulatory genes such as *ctrA*, *ccrM*, *cpdR1*, *divJ*, *divK*, *gcrA* and *pleC*, have been identified in *S. meliloti* (12–16). In addition, the hybrid histidine kinase CbrA is linked to the CtrA regulatory system, which is an important regulator of cell division in *S. meliloti* (17, 18). However, it is still unclear whether these regulatory proteins conduct environmental nutrition signals (e.g., ammonia nitrogen) and whether they play a regulatory role in the symbiotic process.

The NtrY/NtrX two-component system, first discovered in *Azorhizobium caulinodans*,regulates nitrogen metabolism under free-living conditions and affects nodulation and nitrogen fixation in the host plant *Sesbania rostrata* (19). Subsequently, *ntrY/ntrX* homologous genes were found in *Rhizobium tropici* to regulate nitrogen metabolism and symbiotic nodulation (20). NtrY/NtrX homologs regulate nitrate uptake in *Azospirillum brasilense* and *Herbaspirillum seropedicae* (21, 22), and this regulatory system has been found to simultaneously control nitrogen metabolism and cellular redox homeostasis in *Rhodobacter capsulatus* (23). Moreover, NtrX is involved in the regulation of cell envelope formation in *R. sphaeroides* (24). In *Brucella abortus*, the histidine kinase NtrY participates in micro-oxygen signaling and nitrogen respiration (25), while the response regulator NtrX controls the expression of respiratory enzymes in *Neisseria gonorrhoeae* (26). Interestingly, the NtrY/NtrX system regulates cell proliferation, amino acid metabolism and CtrA degradation in *Ehrlichia chaffeensis* (27). Finally, NtrX is required for the survival of *C. crescentus* cells and its expression is induced by low pH (28). These findings show that NtrY/NtrX appears to be a regulatory system for nitrogen metabolism, which may be involved in the regulation of cell division.

NtrX is an NtrC family response regulator protein, which consists of a receiver (REC) domain and a DNA-binding domain (29, 30). X-ray crystal diffraction results indicate that the NtrX protein of *B. abortus* can form a dimer; the REC domain is composed of 5 α-helices and 5 β-sheets; the DNA-binding domain contains an HTH motif, which includes 4 α-helices. The three-dimensional structure of the C-terminus has not been resolved (30). *In vitro* experiments demonstrated that the NtrX protein of *B. abortus* can recognize and bind to the palindromic DNA sequence (**CAA**N_3-5_**TTG**) in the *ntrY* promoter region via the HTH motif to regulate gene transcription (29, 30). In *S. meliloti* 1021, our previous work found that NtrX protein can regulate bacterial growth and proliferation, flagellar synthesis and motility, succinoglycan production, and symbiotic nodulation and nitrogen fixation with the host plant alfalfa (31, 32). In the present study, we investigated the control mechanism by which NtrX regulates *S. meliloti* cell division at the transcriptional level.

## RESULTS

### Defects of cell division resulting from *ntrX* mutation in *S. meliloti*

We previously constructed a plasmid insertion mutant of the *ntrX* gene in *S. meliloti* 1021, called as SmLL1 (31). This mutant grew slowly in LB/MC medium compared to wild-type Sm1021 (31). According to the determined growth curve, the doubling time of bacterial cell proliferation was calculated to be 180 mins for SmLL1 compared to 160 mins for the wild-type strain (33). Microscopic observation revealed that 5% to 10% of SmLL1 cells grown in the LB/MC broth up to the logarithmic phase exhibited morphological abnormalities (such as cell elongation, Y-shaped or V-shaped), whereas Sm1021 had almost no abnormally shaped cells (Fig. 1A-B). To determine whether the appearance of abnormally shaped cells of the SmLL1 strain is associated with the synthesis and segregation of genomic DNA, we synchronized the *S. meliloti* cells, inoculated them in LB/MC broth, grew them for 180 mins, and then harvested the cells for flow cytometric analysis. The results showed that most of the Sm1021 cells were haploids, only a few diploids, whereas the most of SmLL cells were diploid (Fig. 1B), indicating a deceleration of replication and segregation of their genomic DNA as compared to the wild type. These observations indicate that the SmLL mutant has cell division defects.

**Fig. 1.**
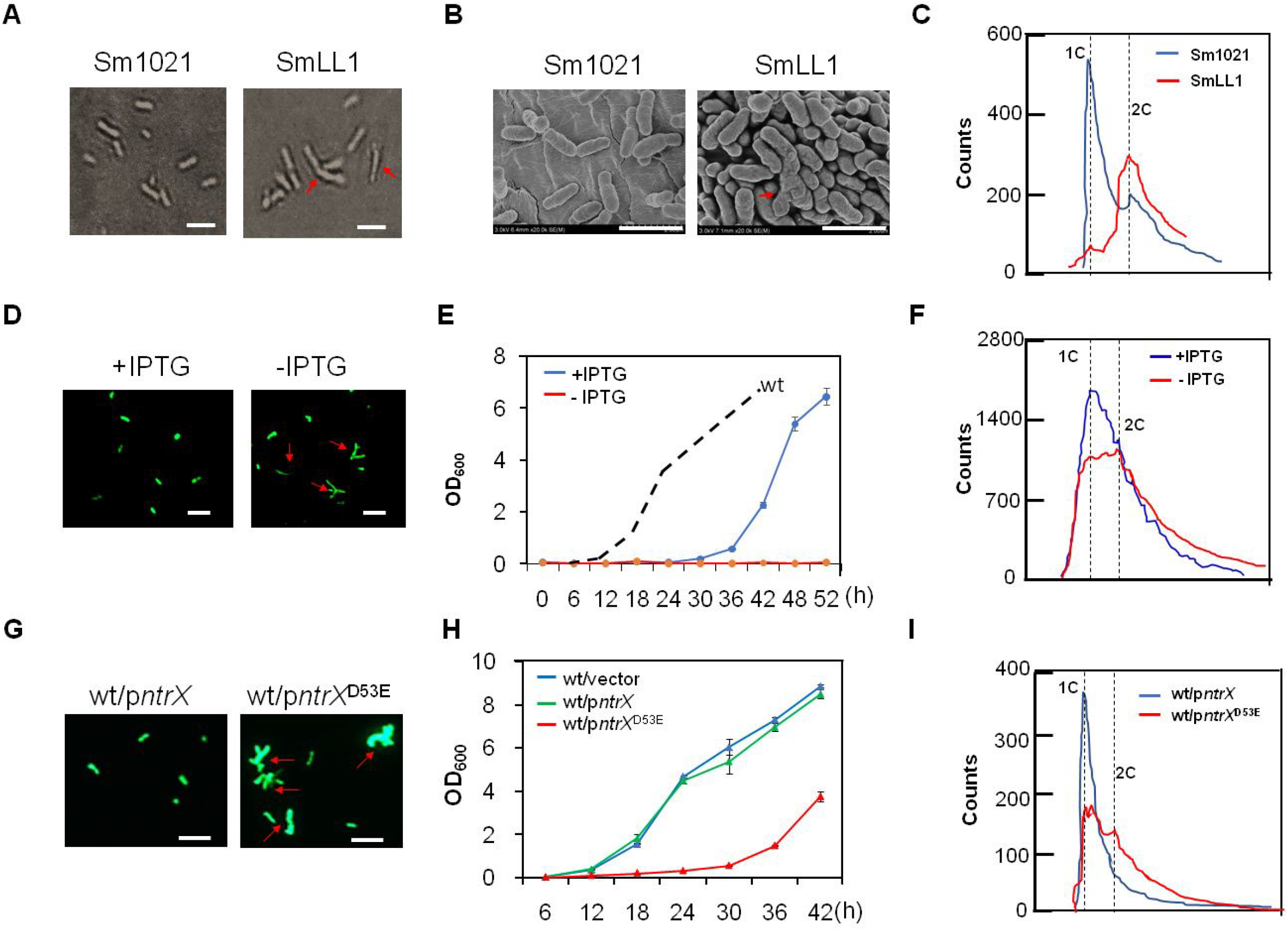
Cell division defects of *S. meliloti ntrX* mutants in LB/MC broth. (**A**, **B**, **D**, **G**) Cell morphology and sizes of *ntrX* mutants under a light, scanning electron or fluorescence microscope. Red arrows, abnormal cells; bars, 2μm. Sm1021 and SmLL1 cells were grown in LB/MC broth to logarithmic phase in **A** and **B.**The depletion cells (Δ*ntrX/pntrX*) carrying pHC60 (35) were grown in LB/MC broth with or without 1 mM IPTG for two hours in **D.** The cells of Sm1021/p*ntrX* and Sm1021/p*ntrX*^D53E^ carrying pHC60 were grown in LB/MC broth containing 1 mM IPTG for two hours. (**C**, **F**, **I**) Genomic DNA content of *ntrX* mutants was determined by flow cytometry. 1C, haploid; 2C, diploid. Synchronized cells of Sm1021, SmLL1, Sm1021/p*ntrX* and Sm1021/p*ntrX*^D53E^ were grown in LB/MC broth for three hours in **C** and **I**. Latter two strains were also incubated with 1mM IPTG in **I**. The depletion cells were grown in LB/MC broth with or without 1 mM IPTG for one hour in **F**. (**E**, **H**) Growth curves of the *ntrX* depletion strain and Sm1021/p*ntrX*^D53E^ in LB/MC broth with 1 mM IPTG. Error bars, ±SD.

Because the deletion of *ntrX* may be fatal, the deletion mutant in *S. meliloti* 1021was not yet successfully screened. Therefore, we constructed a depletion strain that the *ntrX* gene on the genome has been deleted, but carries an IPTG inducible-expression *ntrX* gene from a plasmid (Δ*ntrX*/p*ntrX*) to verify the above results. Optical microscopic observation showed that more than 30% of the *ntrX* depleted cells in LB/MC broth without IPTG displayed abnormal shapes (such as elongation and T-shaped), while in LB/MC broth with IPTG, almost no abnormal cells were observed (Fig. 1D). The depletion strain barely proliferated in LB/MC broth without IPTG, whereas it duplicated slowly with IPTG induction (Figure 1E), indicating that *ntrX* gene expression is required for the cell division of *S. meliloti*. Flow cytometric analysis showed that three peaks were detected in the depletion cells, including haploid and diploid in LB/MC broth without IPTG induction (Fig.1F). After the one-hour induction of IPTG, the peaks were similar to the wild-type cells (Fig. 1F), indicating that genomic DNA replication and segregation of *S. meliloti* requires the expression of the *ntrX* gene.

NtrX, as a regulator of nitrogen metabolism, is composed of a REC domain and a DNA-binding domain (30). The phosphorylated NtrX has been reported in *C. crescentus* and *B. abortus* (28, 30), the putative phosphorylation site is predicted as the conserved 53^rd^ aspartate residue (D53) on the REC domain (Fig. 4A-B). If the NtrX protein is indeed involved in the regulation of *S. meliloti* cell division, as described above, then the mutation of the conserved D53 residue would affect its regulatory function. To test this hypothesis, we tried to construct the substitutions of D53 (replaced by A, N or E) of NtrX from the genome of *S. meliloti* 1021, but were unable successfully to screen the mutants. As a result, we cloned the mutation gene into the expression vector pSRK-Gm (34) and then introduced the recombinant plasmid into Sm1021. On the LB/MC/IPTG plate, we found that the strain expressing NtrX^D53A^ or NtrX^D53N^ almost did not form visible colonies with IPTG induction; however, the strain expressing NtrX^D53E^ or NtrX formed many colonies in the same condition (Fig. S1). GFP-labeled *S. meliloti* cells (35) cultured in LB/MC/IPTG broth up to the logarithmic phase were observed under a fluorescence microscope, and more than 20% of Sm1021/p*ntrX^D53E^* cells had abnormal morphology, while Sm1021/p*ntrX* cells were almost normal (Fig. 1G). The growth curve determination also showed that the growth of Sm1021/p*ntrX^D53E^* in LB/MC/IPTG broth was apparently slower than that of Sm1021/p*ntrX* (Fig. 1G). Synchronized *S. meliloti* cells were subcultured into LB/MC/IPTG broth and grown for 180 mins for flow cytometric analysis. The results showed that only one sharp peak (haploid) was found in Sm1021/p*ntrX* cells, whereas three peaks were detected in Sm1021/p*ntrX^D53E^* cells, including haploid and diploid (Fig. 1H). These results suggest that the D53 phosphorylation of the NtrX protein is required for the regulation of cell division of *S. meliloti*.

### Transcription of cell cycle regulated genes under the regulation of NtrX in *S. meliloti*

Since NtrX is involved in controlling cell division of *S. meliloti*, does it regulate the transcription of cell cycle regulatory genes associated with CtrA system? To test this possibility, we performed a preliminary RNA-Seq analysis between Sm1021 and SmLL1 cells. The results indicated that many genes of cell cycle regulation such as *chpT*, *sciP*, *dnaA*, *ftsZ1*, *ccrM*, *podJ1*, *cckA*, *cbrA*, *pleD*, *divK*, *cpdR1*, *mucR* and *clpP* were differentially expressed in the mutant strain compared with the wild-type strain (Fig. S2 and Table S1), suggesting that transcription of many cell cycle regulatory genes is regulated by NtrX.

To confirm the above results in detail, we applied quantitative RT-PCR to analyze the transcript levels of cell cycle regulatory genes in *S. meliloti* cells. The synchronized Sm1021 and SmLL1 cells were subcultured into LB/MC broth for shaking incubation, and then total RNA was extracted from cells grown for every half an hour. The qRT-PCR results showed that the transcript level of the *ntrX* gene increased first in Sm1021, then decreased, and reached the maximum value in the cells cultured for 90 min, displaying a trend of cyclical changes, while the *ntrY* gene cyclical transcription trend was not obvious (Fig. 2A). Known cell cycle regulatory genes, such as *ctrA*, *gcrA* and *dnaA*, also exhibited a cyclical transcription trend (Fig. 2A and S3A). Compared to the wild-type cells, transcript levels of the *ntrX* gene were significantly low in the SmLL1 cells grown at different times, but the cyclical trend was unchanged, and cell cycle regulatory genes such as *dnaA*, *ftsZ1*, *pleC*, *chpT* and *cpdR1* showed similar down-regulation trend (Fig. 2A and S3A). Contrary to these results, *ctrA* was gradually up-regulated in the SmLL1 cells, and *gcrA*, *ccrM* and *ntrY* were significantly upregulated at the same time (Fig. 2A and S3A). These findings suggest that the NtrX protein may repress the transcription of genes such as *ctrA* and *gcrA* and activate the transcription of genes such as *dnaA* and *ftsZ1*.

**Fig. 2.**
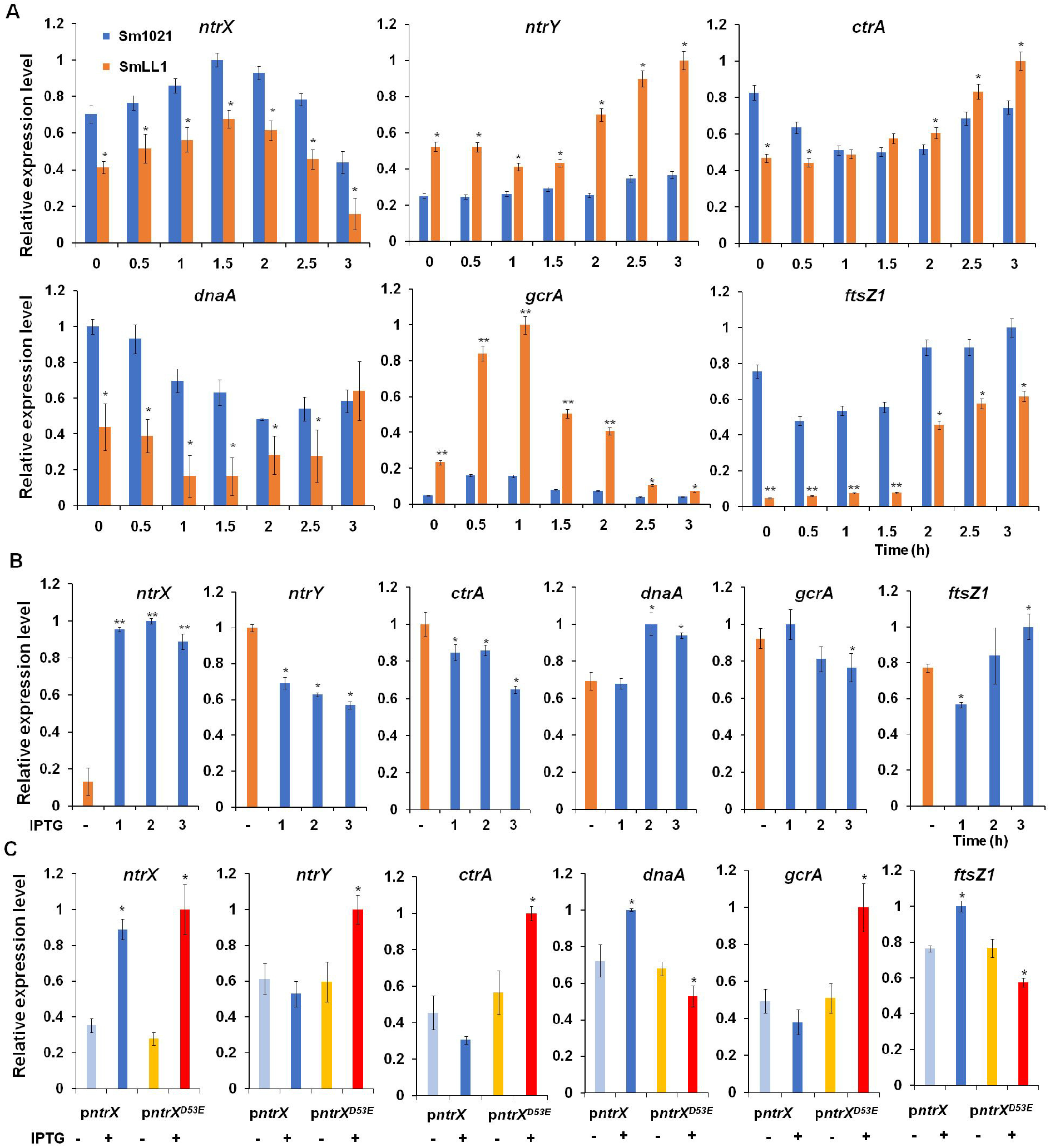
Differential expression of cell cycle regulatory genes in *ntrX* mutants evaluated by qRT-PCR. Synchronized cells of Sm1021 and SmLL1 were grown in LB/MC broth for half to three hours in **A**. The depletion cells were grown in LB/MC broth with 1 mM IPTG for one to three hours, while the cells were incubated without IPTG for one hour as a control in **B**. Synchronized cells of Sm1021/p*ntrX* and Sm1021/p*ntrX*^D53E^ were grown in LB/MC broth with 1 mM IPTG for three hours in **C**. Error bars, ±SD. The student t-test was used for significance analysis. *, P<0.05; **, P<0.001.

We analyzed the transcripts of cell cycle regulatory genes in cells of the depletion strain to verify the above results. The qRT-PCR results showed that depleted cells cultured in LB/MC broth without IPTG had extremely low levels of *ntrX* transcripts, while transcripts of most cell cycle regulatory genes were high-level detected (Fig. 2B and S3B). After culturing the depleted cells in LB/MC broth with 1 mM IPTG for 1 h, numerous *ntrX* gene transcripts were detected (Fig. 2B and S3B). The transcript levels of cell cycle regulatory genes were significantly altered in depleted cells cultured in the broth with IPTG for 2 or 3 h compared to the cells cultured in the broth without IPTG: the transcription of *ctrA, gcrA* and *ccrM* was down-regulated; the transcription of *dnaA*, *ftsZ1*, *pleC*, *chpT* and *cpdR1* was up-regulated (Fig. 2B and S3B). These results further confirm that the NtrX protein represses the transcription of genes such as *ctrA* and *gcrA* and simultaneously activates the transcription of genes such as *dnaA* and *ftsZ1*.

We analyzed the transcript levels of major cell cycle regulatory genes in Sm1021/p*ntrX^D53E^* and Sm1021/p*ntrX* cells to determine whether the conserved D53 residue on NtrX is essential for transcriptional regulation. The qRT-PCR results showed that transcripts of the *ntrX* gene were significantly increased in cells cultured in LB/MC broth containing IPTG for 2 h compared to the cells without IPTG treatment; meanwhile, the transcripts of *ctrA*, *gcrA* and *ntrY* were significantly reduced or tended to decrease, while the transcripts of *dnaA* and *ftsZ1* genes were significantly increased (Fig. 2C). Contrary to the above results, as transcripts of the *ntrX*^D53E^ gene increased significantly under IPTG induction, transcripts of *ctrA*, *gcrA*,and *ntrY* also increased significantly, while transcripts of *dnaA* and *ftsZ1* genes decreased significantly (Fig. 2C). These results further confirm that the NtrX protein represses the transcription of genes such as *ctrA* and *gcrA* and activates the transcription of genes such as *dnaA* and *ftsZ1*, and that this regulation depends on the D53 residue on NtrX.

To determine whether the expression of the CtrA system genes is regulated by NtrX in heterogeneous cells, the promoter-*uidA* fusions were co-transformed with p*ntrX* or the empty vector(pSRK-Gm) into *E. coli* DH5α, respectively. Quantitative analysis of GUS activities showed that the weaker activities of the promoter of *ctrA* or *gcrA* in the cells carrying *pntrX* than those cells with pSRK-Gm were observed (Fig. S3C). In contrast, the activity of the *dnaA* promoter is apparently elevated in the cells co-expressing *ntrX* compared with those cells carrying pSRK-Gm (Fig. S3C). These data supported the conclusion that NtrX negatively controls transcription of *ctrA* and *gcrA*, but positively regulates transcription of *dnaA*.

### Negative regulation of protein levels of CtrA and GcrA by NtrX in *S. meliloti*

To determine whether protein levels of key cell cycle regulators are affected by the *ntrX* mutation, we first expressed His-tagged NtrX, CtrA and GcrA proteins in *E. coli*. After purification by nickel columns, rabbit polyclonal antibodies were prepared for immunoblotting assays (36). The results showed a varying trend of increasing first and then decreasing NtrX protein levels and a maximum occurring in the synchronized cells subcultured for 1.5 h (Fig. 3A). Unlike Sm1021, the total NtrX protein level in SmLL1 cells was apparently reduced and tended to increase gradually at different growth times (Fig. 3A). Contrary to the NtrX protein, the change trend of CtrA and GcrA protein levels in Sm1021 first decreased and then increased. The levels of these two proteins were apparently increased in SmLL1 cells compared to Sm1021 cells, (Fig. 3A). These results indicate that SmLL1 is a down-regulated mutant of *ntrX* and that NtrX protein levels are negatively correlated with CtrA and GcrA proteins.

**Fig. 3.**
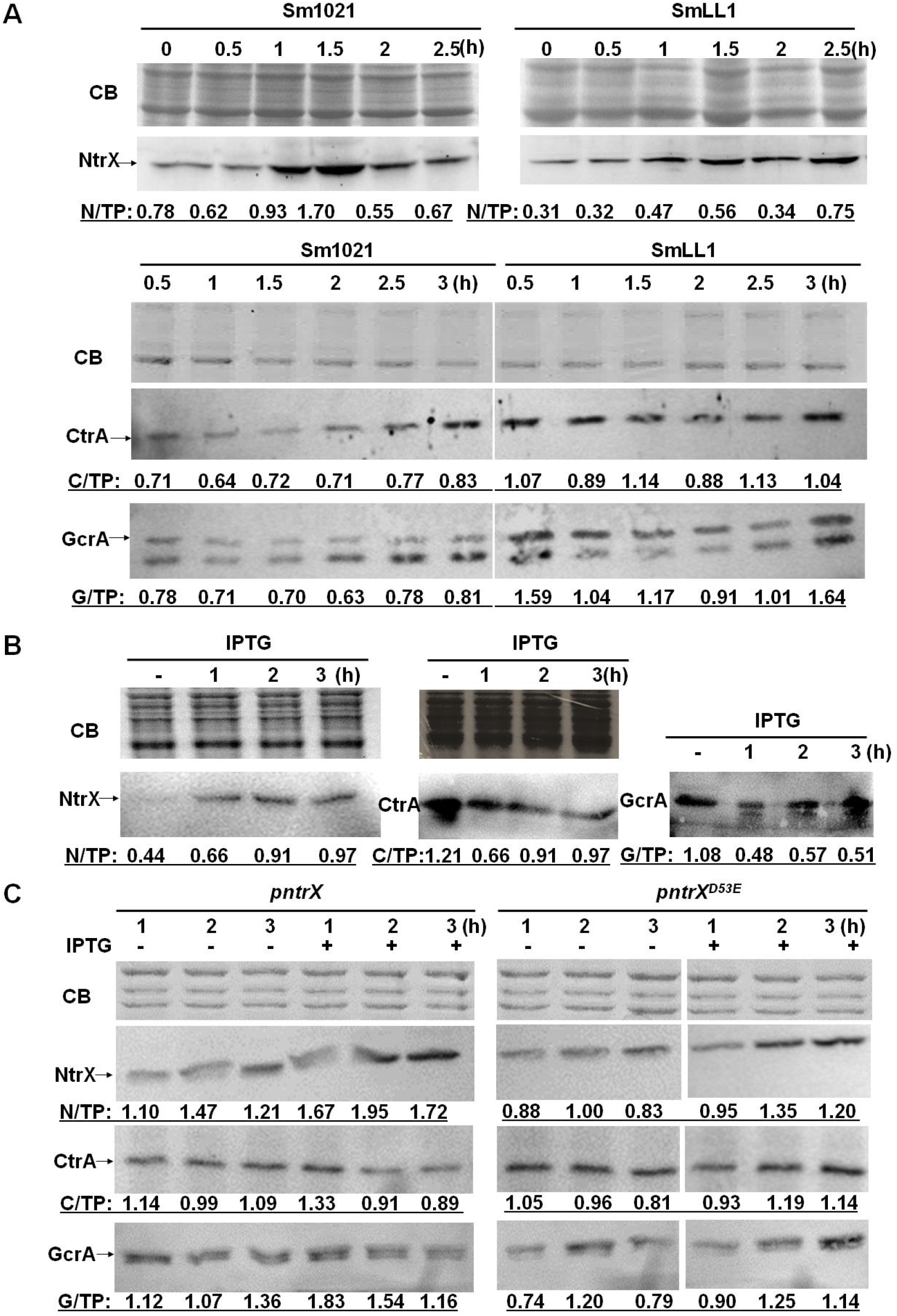
Protein levels of NtrX, CtrA and GcrA in the *ntrX* mutant as evaluated by Western blotting. Synchronized cells of Sm1021 and SmLL1 were grown in LB/MC broth for half to three hours in **A**. The depletion cells were grown in LB/MC broth containing 1 mM IPTG for one to three hours, while the cells were incubated without IPTG for one hour as a control in **B**. Synchronized cells of Sm1021/p*ntrX* and Sm1021/p*ntrX*^D53E^ were grown in LB/MC broth containing 1 mM IPTG for one to three hours in **C**. CB, total proteins stained by Coomassie brilliant blue. N/TP, C/TP or G/TP, the blot intensity of NtrX, CtrA or GcrA (the larger one) divided the intensity of the strongest band of the total proteins stained by Coomassie brilliant blue. The intensity data were collected using Image J (42).

We evaluated the NtrX protein level in cells of the depletion strain grown in LB/MC broth by immunoblotting and found that cells cultured in the broth containing IPTG for 1 to 3 h high-level expressed NtrX protein (Fig. 3B). CtrA and GcrA proteins were high-level expressed in cells cultured in LB/MC broth without IPTG, whereas their levels were apparently reduced in cells cultured in broth containing IPTG for 1 to 3 h (Fig. 3B). These results also prove that NtrX protein levels are negatively correlated with CtrA and GcrA proteins.

To determine whether the D53 residue on the NtrX protein affects the protein levels of CtrA and GcrA, we performed immunoblot assays of lysates from Sm1021/p*ntrX*^D53E^ and Sm1021/p*ntrX* cells. The results showed that the protein levels of NtrX and NtrX^D53E^ increased apparently when cultured in LB/MC broth containing IPTG for 2-3 h (Fig. 3C). Under the same culture conditions, the protein levels of CtrA and GcrA were reduced somewhat in Sm1021/p*ntrX* cells, while they were elevated to some extent in Sm1021/p*ntrX*^D53E^ cells (Fig. 3C). These results reaffirm that NtrX protein negatively regulates the expression of CtrA and GcrA in the dependent manner of the D53 residue.

### The 53^rd^ aspartate residue as a phosphorylation site of *S. meliloti* NtrX

The homologous NtrX proteins in α-proteobacteria are composed of REC and DNA binding domains. The three-dimensional structure of the NtrX protein from *B. abortus* has been partially resolved (29, 30). Using this as a template, we reconstructed the 3D structure of the NtrX protein from *S. meliloti* and found that there were 5 α-helices and 5 β-sheets connected by loops in the REC domain (Figure 4A-B). The conserved D53 is located at the end of the third β-sheet and predicted as a phosphorylated residue by PFAM.

**Fig. 4.**
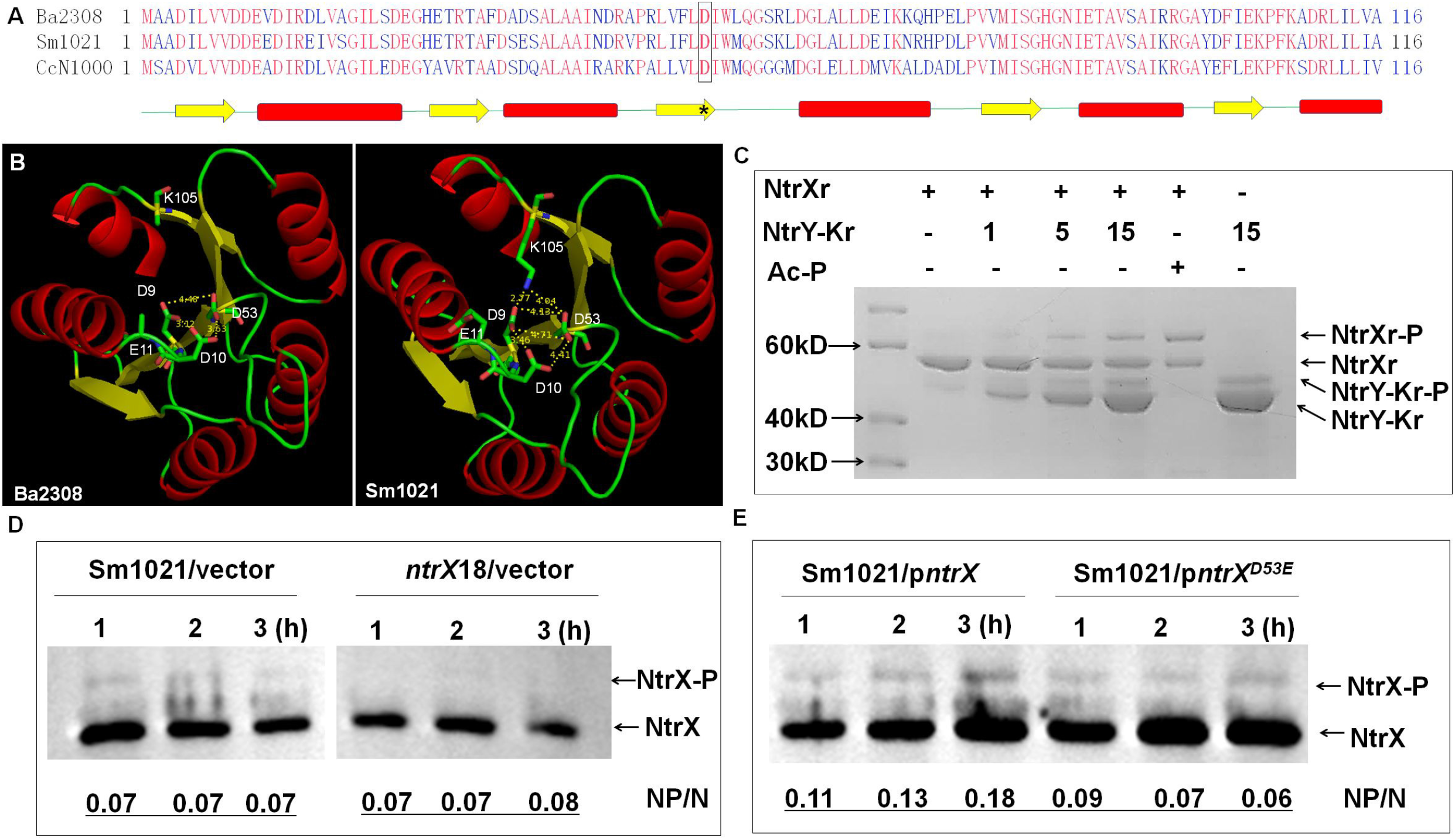
Phosphorylation of the 53^rd^ aspartate residue in the NtrX protein. (**A**) Alignment of NtrX receiver domain from three bacterial species. The amino acid sequence of each protein was obtained from NCBI. Secondary structures of the receiver domain are shown as green lines (loops), yellow arrows (β-sheets) and red bars (α-helixes). Both the aspartate residue in the box and the asterisk represent the predicted phosphorylation site from Pfam. Ba2308, *B. abortus* bv. 1 str. 2308; Sm1021, *S. meliloti* 1021; CcN1000, *C. crescentus* N1000. (**B**) 3D structure of the NtrX receiver domain of *S. meliloti* NtrX. It was reconstructed using *B. abortus* homolog protein (PDB: 4d6y) as a template in Swiss-Model. Possible electrostatic interactions associated with the 53^rd^ aspartate residue are labeled via Pymol. (**C**) *In vitro* NtrX phosphorylation catalyzed by the NtrY kinase domain. NtrY-Kr, His-NtrY kinase domain fusion protein (1, 3 and 10 μg); NtrXr-P, phosphorylated NtrXr (10 μg); Ac-Pi, acetyl phosphate (2 mM). (**D-E**) *In vivo* phosphorylation of *S. meliloti* NtrX. Phosphorylated NtrX proteins from *S. meliloti* cells were separated by Phos-Tag gel and detected by Western blotting of anti-NtrX polyclonal antibodies. *S. meliloti* 1021 cells carrying p*ntrX* or p*ntrX*^D53E^ were grown in LB/MC broth with 1mM IPTG for 1 to 3 h. ~ 1 μg of total protein was loaded into each well. NP/N, the intensity of phosphorylated proteins divided the intensity of non-phosphorylated proteins. The intensity data were collected using Image J (42).

From the report of *B. abortus*, the NtrY histidine kinase can phosphorylate NtrX *in vitro* (29, 30). We expressed and purified the NtrY kinase domain and NtrX protein of *S. meliloti* in *E. coli* for *in vitro* phosphorylation assays. Through Phos-Tag Gel assays, we found that the NtrY kinase domain was autophosphorylated, and phosphorylated the NtrX protein *in vitro* (Fig. 4C). After mutating the 53^rd^ aspartate to glutamate, phosphorylated NtrX protein was not detected by treatment of acetyl phosphate (data not shown). To further determine whether NtrX is phosphorylated *in vivo*, western blotting assays were performed using anit-NtrX antibodies after separating phosphorylated proteins of *S. meliloti* cells by Phos-Tag Gel. The results showed that more phosphorylated NtrX proteins were detected in Sm1021 cells than those in SmLL1 cells as the same as the unphosphorylated protein (Fig. 4D). To further verify that the D53 residue is the phosphorylation site of the NtrX protein, we applied the same method to analyze the phosphorylated NtrX protein level of Sm1021/p*ntrX*^D53E^ cells cultured in LB/MC/IPTG broth. The results showed that the ratio of phosphorylated NtrX protein compared to unphosphorylated protein in Sm1021/p*ntrX* cells tended to increase, whereas the ratio in Sm1021/p*ntrX*^D53E^ cells tended to decrease (Fig. 4D). These results reveal that the D53 residue of NtrX is phosphorylated in *S. meliloti* cells.

### Direct binding of phosphorylated NtrX protein to the promoter DNA of key cell cycle regulatory genes

To determine whether the NtrX protein of *S. meliloti* directly regulates the expression of cell cycle regulatory genes, we used anti-NtrX polyclonal antibodies with high specificity (Fig. S4) to perform chromatin immunoprecipitation experiments. Sequencing results showed that a total of 82 DNA fragments were specifically precipitated from Sm1021 cells, 60 of which were derived from the chromosome, while the other 22 fragments originated from the plasmids SymA and SymB (Fig. 5A). After sequence analysis in detail, we found that the promoter DNA fragments of cell cycle regulatory genes such as *ctrA*, *dnaA*, *mucR* and *cpdR1* were specifically enriched (Fig. 5B and Table S2). Due to of the recognition sites (CAAN_3-5_TTG) of NtrX on the *ntrY* gene promoter reported in *B. abortus* (30), we searched them in the precipitated DNA fragments, and found that nine of possible motifs are located in the promoter regions of cell cycle regulatory genes such as *ctrA*, *danA*, *gcrA* and *ftsZ1*(Fig. S5). To verify the ChIP-Seq results, we applied quantitative PCR to evaluate the level of genomic DNA fragments precipitated by anti-NtrX polyclonal antibodies. The results showed that the promoter regions of *ctrA*, *dnaA*, *gcrA* and *ftsZ1* genes were enriched to different degrees (Fig. 5C), indicating that the NtrX protein in Sm1021 cells can interact directly with the promoter regions of the aforementioned cell cycle regulatory genes.

**Fig. 5.**
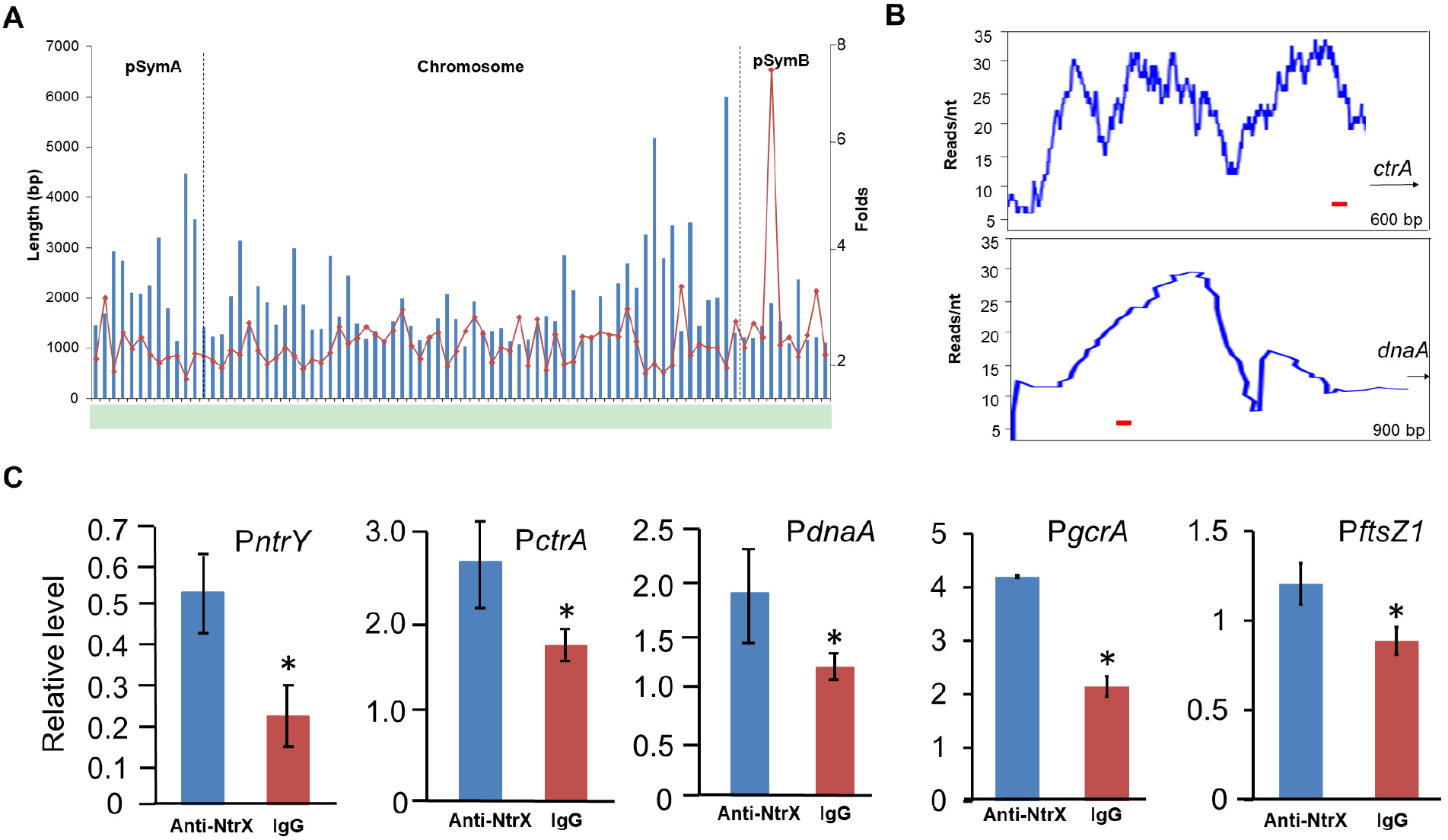
NtrX binding to the promoter DNA of key cell cycle regulatory genes *in vivo*. (**A**) Genome-wide distribution of DNA fragments precipitated by anti-NtrX antibodies through a ChIP-Seq assay. (**B**) Peak maps showing the promoter fragments of *ctrA* and *dnaA* in the ChIP-Seq assay. Red bars, the putative NtrX recognition site, CAAN2-5TTG. (**C**) Enrichment of DNA fragments containing *ctrA*, *gcrA* and *dnaA* promoters determined by ChIP-qPCR. IgG, *S. meliloti* lysate treated by IgG as a negative control. Error bars, ±SD. The student t-test was used for significance analysis. *, P<0.05.

In Sm1021, the promoter DNA of the *ntrY* gene can directly interact with the NtrX protein (Fig. 5C), which is similar to the report in *B. abortus* (30). To further confirm the above results, we synthesized a biotin-labeled probe of *ntrY* promoter DNA (containing two **CAA**N_3-5_**TTG** motifs: **CAA**CACCG**TTG** and **CAA**TGCG**TTG**) for gel retardation assays. The results showed that phosphorylated NtrX specifically bound to it, forming two protein-DNA complexes (Fig. 6A). To determine whether the D53 phosphorylation of the NtrX protein is involved in the protein-DNA binding reaction, we replaced the phosphorylated NtrX protein with the NtrX^D53E^ protein. The gel retardation results showed almost no protein-DNA complex formation (Fig. 6A), suggesting that the phosphorylation of D53 is essential for the binding of NtrX to the *ntrY* promoter region. The same method was used to analyze the binding ability between the phosphorylated NtrX protein and the biotin-labeled probe of *dnaA* promoter DNA (containing the **CAA**AACCC**TTG** motif) and found that they bound specifically to form a protein-DNA complex (Fig. 6B). We mutated the **CAA**AACCC**TTG** motif of the DNA probe to **CGG**AACCC**CCG**, and found that the mutant probe virtually did not bind to the phosphorylated NtrX protein (Fig. 6B), suggesting that the base composition of the recognition site is important for NtrX binding reaction. We also used gel retardation assays to confirm whether the phosphorylated NtrX protein can specifically bind to biotin-labeled probes of *ctrA*, *gcrA* and *ftsZ1* promoter DNA (each containing a **CAA**N_3-5_**TTG** motif: **CAA**CC**TTG**, **CAA**ACC**TTG** and **CAA**TGGC**TG**), and found that at least one protein-DNA complex was formed, respectively (Fig. 6C-E). These results indicate that the phosphorylated NtrX protein can bind specifically to the promoter regions of *ctrA*, *gcrA*, *dnaA* and *ftsZ1 in vitro*.

**Fig. 6.**
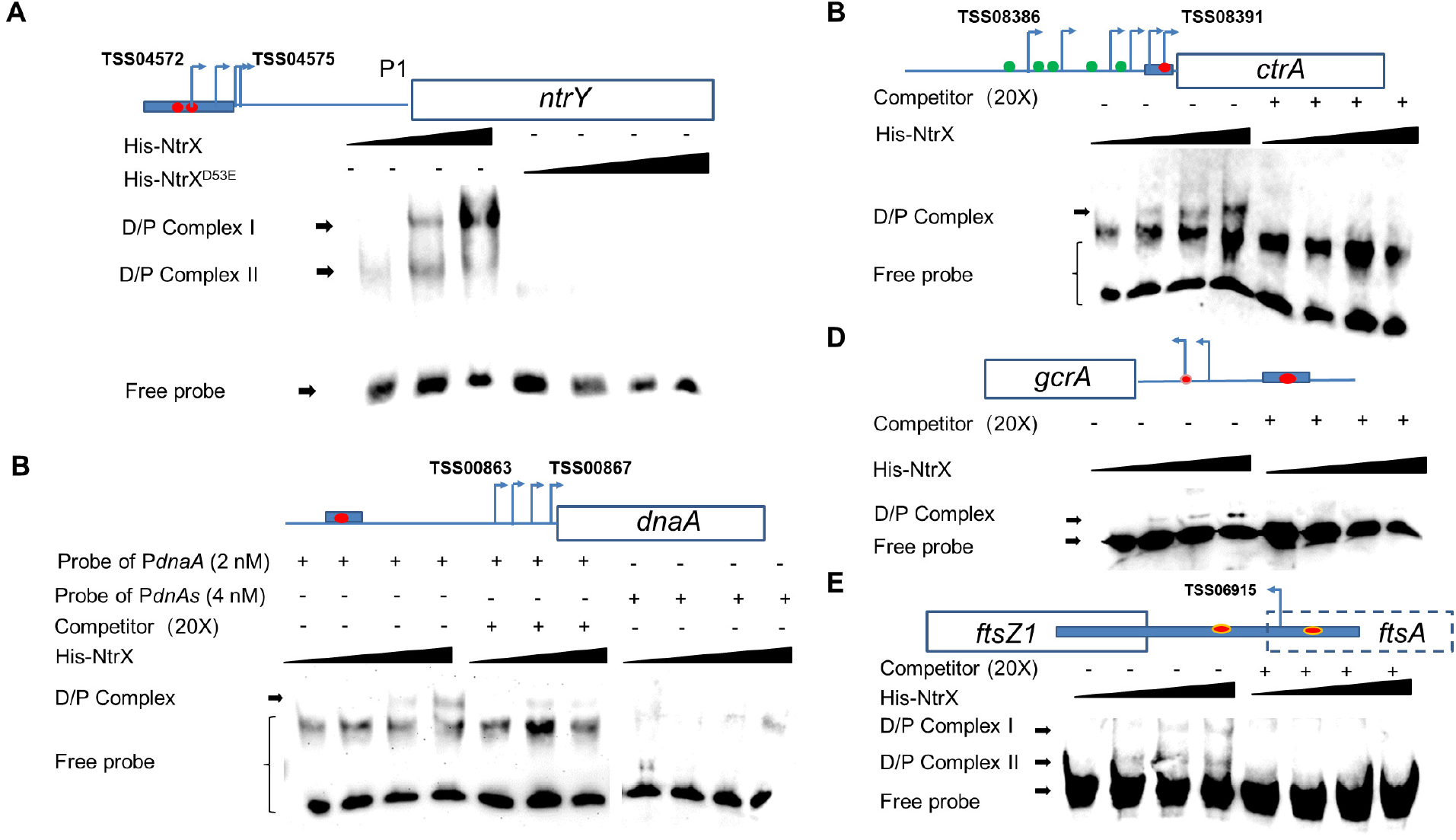
Phosphorylated NtrX proteins binding to the promoter DNA of *ntrY* (**A**), *ctrA* (**B**), *dnaA* (**C**), *gcrA* (**D**) and *ftsZ1*(**E**) *in vitro*. His-NtrX^D53E^, the His-NtrX fusion protein containing a substitution of D53E in (**A**). Probe *PdnaAs* is the DNA probe *PdnaA* that C**AA**AACCC**TT**G was replaced by C**GG**AACCC**CC**G in (**B**). D/P complex, DNA-protein complex; competitor, the DNA probe without biotin labeling. 0, 3, 6 and 15 ng His-NtrX proteins mixed with each probe (2 nM), respectively. TSS, transcriptional start sites from the literature (**37**). Blue bars, probes for EMSA; red balls, the putative recognition site of NtrX, CAAN2-5TTG; green balls, the binding sites of CtrA (12, 13).

## DISCUSSION

In symbiotic nitrogen-fixing bacteria, rhizobia, the level of combined nitrogen as a signal not only regulates the expression of nitrogen-fixing genes, but also can affect cell growth and division. However, the molecular mechanism by which combined nitrogen levels regulate bacterial cell division is unclear. This work first revealed in *S. meliloti* that the nitrogen metabolism regulator NtrX directly regulates the transcription of cell cycle regulatory genes such as *ctrA*, *gcrA*, *dnaA* and *ftsZ1* by specifically interacting with the promoter regions, to promote cell division (Fig. 7), which provides a preliminary answer to the above question.

**Fig. 7.**
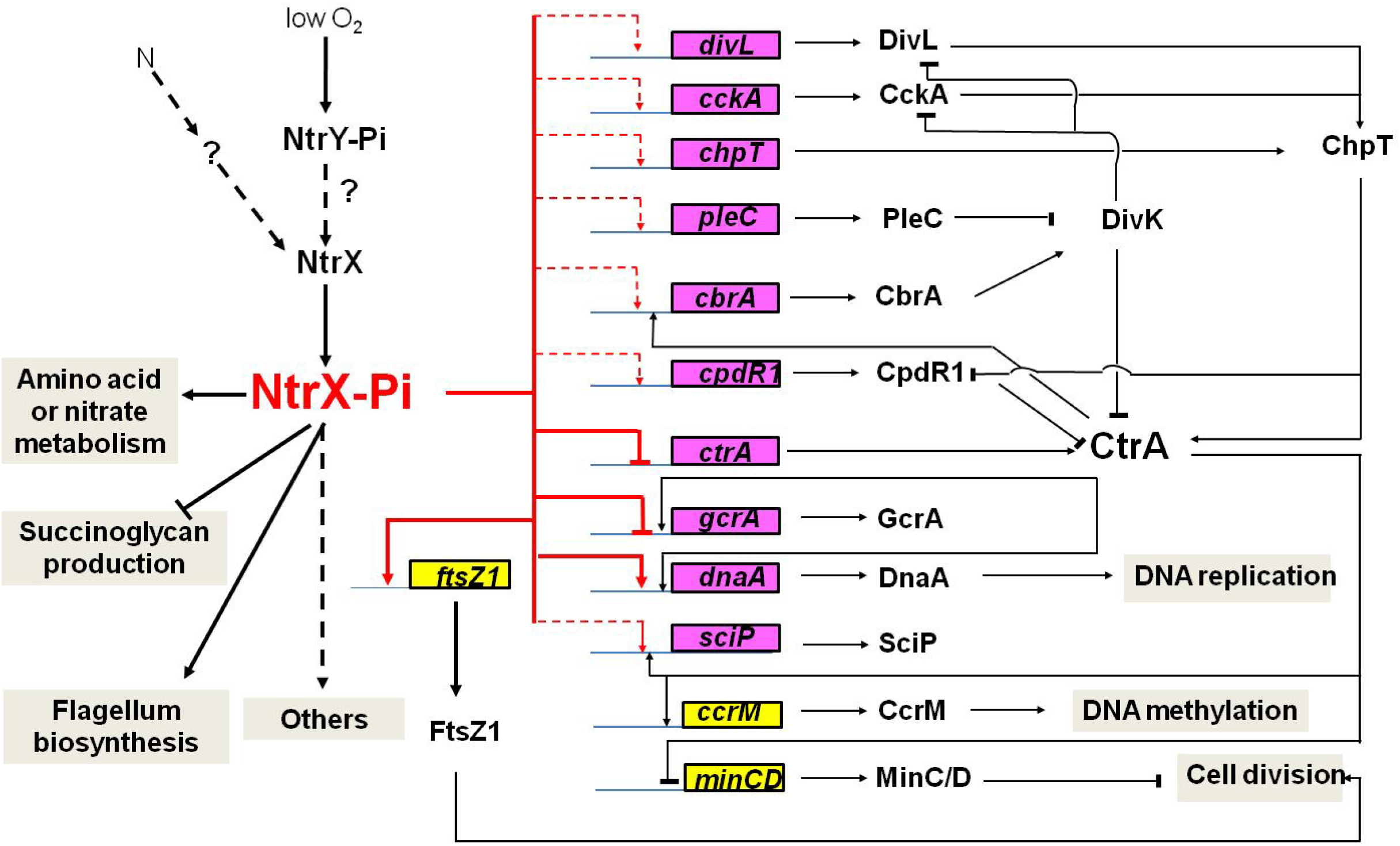
An NtrX-mediated transcriptional control system for cell cycle progression of *S. meliloti*

NtrX is a bacterial cell cycle regulator. Previous studies have suggested that NtrX is a regulator of nitrogen metabolism in bacterial cells because its mutants affect the utilization of nitrogen sources and NtrX is able to regulate amino acid metabolism and nitrogen oxidation (19–22, 25, 27). Decreased utilization of nitrogen sources would inevitably lead to weakened nucleic acid and protein synthesis, which would subsequently suppress the growth and proliferation of bacterial cells. This is one explanation to the effect of *ntrX* mutations on bacterial cell division. In *E. chaffeensis* cells, NtrX affected the stability of the CtrA protein through a post-translational mechanism (27), indicating that NtrX may act directly on the cell cycle regulatory system to regulate cell division. This work was carried out using *S. meliloti* as the study material and reveals the transcriptional control mechanism of the cell cycle regulatory genes mediated by the NtrX protein for the first time. This conclusion is supported by multiple experimental evidences: 1) three sets of *ntrX* gene mutation materials are defective in bacterial growth, cell morphology and genomic DNA synthesis (Fig. 1); 2) the transcript levels of cell cycle regulatory genes such as *ctrA*, *gcrA*, *dnaA* and *ftsZ1* are differentially altered in *ntrX* mutants (Fig. 2; 31; 3) the protein levels of CtrA and GcrA are correspondingly altered in the mutant and they were negatively correlated with the level of NtrX protein (Fig. 3); 4) the phosphorylated NtrX protein binds directly to the promoter regions of *ctrA*, *gcrA*, *dnaA* and *ftsZ1* (Fig. 5–6).

Cell division defects showed a little difference for three sets of *ntrX* mutants. For example, the fewest cells of abnormal shapes were observed for SmLL1 cells, the most cells were found for the depletion cells (Fig. 1A-B, D, G). In fact, the depletion cells are easy to die in LB/MC medium without IPTG addition. Even in LB/MC broth containing different concentrations of IPTG, it still grew slower than the wild-type strain, Sm1021 (Fig. 1E). Irregular cell shapes of the *ntrX* mutant may be associated with genomic DNA content. We noticed that the flow cytometric peak of haploid cells is very sharp for Sm1021 or Sm1021/p*ntrX*, but it was not for SmLL1, the depletion strain and Sm1021/p*ntrX*^D53E^ (Fig. 1C, F, I).

Differential expression of cell cycle regulatory genes exhibited some difference for three sets of *ntrX* mutation materials. First, qRT-PCR data showed larger expression differentials between SmLL1 and Sm1021 than those of the depletion strain (with and without IPTG induction) and the Sm1021/p*ntrX*^D53E^ strain (compared with Sm1021/p*ntrX*, Fig. 2 and S3A-B). It may be result from the different cell cycle status of tested cells. Secondly, preliminary transcriptomic data showed more cell cycle regulatory genes differentially expressed between the synchronized cells of SmLL1 and Sm1021 than those cells subcultured in LB/MC broth (Fig. S2). Expression differentials of these genes from the cells subcultured in LB/MC broth were further determined by qRT-PCR (Fig. 2A and Fig. S3A), since this method is more sensitive for mRNA level analysis than RNA-seq from our experience.

Transcriptional regulation of *ctrA*, *gcrA* and *dnaA* mediated by NtrX is confident, though the expression differentials are varied from different materials or detected by different methods (Fig. 2 and S2). This conclusion was supported by heterogeneous expression and western blotting results (Fig. S3C and 3). Moreover, the expression results coincide with data of interactions between of NtrX and promoter regions of *ctrA*, *gcrA* and *dnaA* (Fig. 5–6). An NtrX homolog may bind to the promoter region of *dnaA* in *R. sphaeroides* based on the published ChIP-seq data (24). NtrX may bind to the recognition sites that contain a transcription start site to prevent transcriptional initiation of *ctrA* and *ntrY* (37) (Fig. 6A, 6C). We noticed that the bands of the complex between NtrX and the *gcrA* probe were relatively weak, which may be associated with the selected probe (Fig. 6D).

NtrX phosphorylation has been reported in *B. abortus* and *C. crescentus*, and it is required for the formation of *ntrY* promoter DNA-NtrX complex in *B. abortus* (28–30). The same result was gained in *S. meliloti* (Fig. 4C and 6A), suggesting that NtrX phosphorylation is conserved in these species. Based on homology and conservativeness of NtrX proteins (Fig. 4A-B), the conserved 53^rd^ aspartate was predicted as the phosphorylation residue. The mutation protein NtrX^D53E^ was neither phosphorylated by acetyl phosphate *in vitro* (data not shown), nor by histidine kinase in *S. meliloti* cells (Fig. 4E), confirming that D53 is the real phosphorylation site. The NtrX^D53E^ may mimic the phosphorylated NtrX protein to retain partial functions, which is completely different from NtrX^D53A^ and NtrX^D53N^ (Fig. S1).

Phosphorylated NtrX can recognize *cis* elements on the promoters of downstream regulated genes in bacterial species (24, 30). These *cis* elements are not completely consistent from different literatures. In *B. abortus*, the NtrX binding sites CAAN_3-5_TTG have been identified in the promoter region of *ntrY* by footprinting (30). In *R. sphaeroides*, the GCAN_9_TGC motifs have been suggested to be NtrX recognition sites by analyzing ChIP-seq data (24). These NtrX recognition sites from above two species share the palindromic sequence CAN_x_TG. We neither found GCAN_9_TGC motifs from the probes specifically binding to NtrX in *S. meliloti*, nor identified them by analyzing our ChIP-seq data (Fig. 5A, 6). Furthermore, at least one CAAN_2-5_TTG motif located in the promoter regions of cell cycle regulatory genes such as *ctrA*, *gcrA*, and *dnaA* were found (Fig. S5). These observations are not only consistent with our footprinting assays of the promoters of *visN* (36) and *dnaA* (data not shown), but also with our EMSA results (Fig. 6, S5). Interestingly, when the NtrX binding site is overlapped with one of transcriptional start sites of *ctrA* and *ntrY*, the expression of these genes is downregulated by NtrX (37) (Fig. 2, 3, 6). The possible explanation is that NtrX binding to the sites prevents transcription initiation of these genes. Although we identified that NtrX binds to the motifs of CAAN_2_TTG of the promoters of *visN* and *ctrA* (36) (Fig. 6C, S5), but we still don’t know why NtrX recognition sites contain the length-varied palindromic sequences.

The upstream kinase of NtrX may be not the cognate kinase NtrY, though the NtrY recombinant kinase from *S. meliloti* and *C. crescentus* phosphorylated NtrX protein *in vitro* (25) (Fig.4C). We noticed that both ORFs of *ntrY* and *ntrX* are overlapped and the repression of *ntrY* gene expression by NtrX in *S. eliloti* (31); Fig. 2). The phenotypes of the *ntrY* deletion mutant did not coincide with the *ntrX* mutant (16). These observations are consistent with the report that NtrY may be the phosphatase of NtrX in *C. crescentus* (38). Moreover, the expression of *ntrY* gene (not *ntrX*) is induced by micro-oxygen in *B. abortus* (25). The primary function of NtrX in bacteria was considered to regulate nitrogen metabolism. The nitrogen limitation signal transduction in bacteria is mainly mediated by the NtrB/NtrC system (39), so it cannot be ruled out that the NtrB/NtrC system can regulate the expression of *ntrX* under nitrogen lacking conditions. Under nitrogen rich conditions, the activity of NtrX may be regulated by an unknown kinase, which may be able to sense the level of combined nitrogen.

## MATERIALS AND METHODS

### Strains and culture medium

*Escherichia coli* DH5α and BL21 were cultured in LB medium at 37 °C. *S. meliloti* (including Sm1021, SmLL1, Δ*ntrX*/p*ntrX* and derivatives) (31) were cultured in LB/MC medium at 28 °C. MOPS-GS broth was utilized for the cell synchronization of *S. meliloti* (33). The following antibiotics were added to the medium: kanamycin (Km), 50 μg/ml; gentamicin (Gm), 10 μg/ml; chloramphenicol (Cm), 30 μg/ml; neomycin (Nm), 200 μg/ml; streptomycin (Sm), 200 μg/ml.

### Recombinant plasmid construction

Primers P*ntrX*1 and P*ntrX*2 bearing *Hind*III and *XbaI* digestion sites were used to amplify the *S. meliloti ntrX* gene (Table S3). The *ntrX* gene fragment was amplified using overlapping PCR primers NMF and NMR with the substitution of aspartate to glutamate, asparagine or alanine (Table S3). Overlapping PCR was performed as described by Wang (31) in 2013. The PCR products were digested with *Hind*III and *Xba*I (Thermo) and ligated with digested pSRK-Gm (34) to obtain the recombinant plasmids p*ntrX*, p*ntrX*^D53E^, p*ntrX*^D53A^ and p*ntrX*^D53N^. Each plasmid was transferred into Sm1021 to gain merodiploids.

After introducing p*ntrX* into SmLL1 by triparental mating with help of MT616, the cells were streaked on LB/MC/Sm agar plates containing 1 mM IPTG and 10% sucrose to screen the *ntrX* depleted cells. The depletion strain (Δ*ntrX*/p*ntrX*) was identified by PCR with the primers of P*ntrYk*1 and P*ntrX*2 (Table S3).

Primers P*ntrY*k1/2, P*ntrX*^D53E^1/2, P*ctrA*1/2, and P*gcrA*1/2 were used to amplify the NtrY kinase domain, *ntrX*^D53E^, *ctrA* and *gcrA*, respectively (Table S3). The DNA fragments amplified by high fidelity PCR (Takara) were digested with the appropriate restriction enzymes, and ligated into pET28b (Sangon) to obtain p*ntrY*k, p*ntrX*^D53E^, p*ctrA* and p*gcrA*, use for recombinant protein purification. The cloned genes on the plasmids were identified by DNA sequencing (Sangon).

Primers P*ctrA*p1/2, P*gcrA*p1/2, P*dnaA*p1/2 were used for amplification the promoter regions of the *ctrA*, *gcrA* and *dnaA*, respectively (Table S3). The PCR fragments were digested by appropriate restriction enzymes and ligated into pRG960 (40) to gain the recombinant plasmids pP*ctrA*, pP*gcrA* and pP*dnaA*.

### Bacterial cell synchronization

De Nisco’s method was used for bacterial cell synchronization (33). *S. meliloti* colonies were selected from an agar plate, inoculated into 5 ml LB/MC broth, and shaken cultured at 28 °C, 250 rpm/min overnight. 100 μl of the bacterial culture was transferred into 100 ml LB/MC broth and shaken cultured overnight until OD_600_ = 0.1-0.15. The cells were collected by centrifugation (6,500 rpm, 5 min, 4 °C), washed twice with sterilized 0.85% NaCl solution, resuspended in MOPS-GS synchronization broth, and shaken cultured for 270 min. After centrifugation, the cells were washed twice with sterilized 0.85% NaCl solution, resuspended in LB/MC broth, and grown at 28 °C.

### RNA extraction, purification and qRT-PCR

The cells from 20 ml of bacterial cultures were collected by centrifugation (6,000 rpm, 5 min, 4°C), and washed twice with DEPC-treated water. RNA extraction was performed using 1 ml of Trizol (Life Technology). Total RNA was treated with genomic DNA Eraser (Takara) to remove any remaining genomic DNA, and then transcribed to cDNA using a PrimeScript RT Reagent Kit (31). The qPCR reaction system included the following: SYBR^®^ Green Real-time PCR Master Mix, 4.75 μl; cDNA or DNA, 0.25 μl; 10 pmol/μl primers, 0.5 μl; ddH_2_O, 4.5 μl. The reaction procedure is as follows: 95°C, 5 min; 95°C, 30 s; 55°C, 30 s; 72°C, 1 kb/min. The selected reference gene was SMc00128. The 2-^ΔΔ^CT method was applied to analyze gene expression levels. All primers are listed in Table S3.

### Chromatin immunoprecipitation (ChIP)

ChIP was performed as described by Pini (4) using rabbit anti-NtrX polyclonal antibodies prepared by Wenyuange, Shanghai (36). In brief, Sm1021 cells (2ml, OD_600_ of 0.8) were cross-linked in 10 mM PBS (pH7.6) containing 1% formaldehyde at room temperature for 10 min, and incubated on ice for 30 min. The cells were washed three times with PBS, treated with lysozyme, sonicated (EpiShear^™^) on ice using 15 bursts of 30 sec (50% duty) at 40% amplitude. Lysates were diluted in 1 mL of ChIP buffer and pre-cleared with 50 μl of protein-A agarose and 80 μg BSA. Anti-NtrX polyclonal antibodies were added to the supernatant (1:1,000 dilution), incubated overnight at 4°C with 50 μl of protein-A agarose beads pre-saturated by BSA, washed with low, high salt and LiCl buffer once and twice with TE buffer. The protein-DNA complexes were eluted using 200 μl freshly prepared elution buffer (1% SDS, 0.1 M NaHCO_3_) supplemented with NaCl to a final concentration of 300 mM, and incubated for 6 h or overnight at 65°C to reverse the crosslinks. DNA was purified by a MinElute kit (QIAGEN) and resuspended in 40 μl of Elution Buffer. DNA sequencing was completed using Illumina HiSeq 2000 in BGI. PCR was performed as the same as above qRT-PCR, and SMc00128 was used as an internal reference to normalize the data.

### Flow cytometry

De Nisco’s flow cytometry protocol was used (33). The cells from 4 ml of *S. meliloti* cultures were collected by centrifugation (6,000 rpm, 5 min, 4°C), washed twice with a 0.85% NaCl solution (stored at 4 °C). 250 μl of cell suspension was mixed with 1 ml of 100% ethanol to fixation. The fixed cells were collected by centrifugation (6000 rpm, 3 min), and incubated in 1 ml of 50 mM sodium citrate buffer containing 4 μg/ml RNase A at 50 °C for 1.5 hours. 1 μl of 10 μM SYTOX Green dye (Sigma) was added to each sample. Each sample was assessed using a MoFlo XDF (Beckman Coulter) flow cytometer, and the results were analyzed by Summit 5.1 software (Beckman Coulter).

### EMSA (electrophoretic mobility shift assay)

EMSA was performed as described by Zeng (36). 30 μl of the purified NtrX protein solution (200 ng/μl) was incubated with 20 μl of 100 mM acetyl phosphate (Sigma) in 50 μl of 2X buffer (100 mM Tris-HCl pH 7.6, 100 mM KCl, 40 mM MgCl_2_) for 1 h at 28 °C. The remaining acetyl phosphate was removed by ultra-filtration (10 KD Amicon Ultra 0.5, Millipore). The protein-DNA binding reaction (20 μl) included 3, 6, 15 ng phosphorylated NtrX protein, 2 or 40 nM DNA probe, 1X binding buffer, 5 mM MgCl_2_, 50 ng/μl poly(dI·dC), 0.05% NP-40, 1% glycerol, and ddH_2_O (up to 20 μl). The mixture was incubated for 30 min at 28 °C, after which 1 μl of loading buffer was added for PAGE. The protein-DNA complexes were transferred onto a nylon membrane (Thermo) and irradiated with a 254 nm UV lamp for 10 min. The protein-DNA complexes were detected using a Light Shift Chemiluminescent EMSA Kit (Thermo). Probes of *ntrY*, *ctrA*, *dnaA*, *gcrA* and *ftsZ1* promoter DNA labeled with biotin were synthesized in Invitrogen, Shanghai, and listed in Table S3.

### NtrX phosphorylation assay and western blotting

The procedure of NtrX phosphorylation assays was modified from Pini (15). 1 mg His-NtrX fusion protein (NtrXr) and His-NtrY kinase domain fusion protein (NtrY-Kr) purified through a Ni^2+^column were used for *in vitro* phosphorylation assays. 2 mM acetyl phosphate (Sigma) was mixed with 300 μg NtrY-Kr in 1 ml of phosphorylation buffer (50 mM Tris-HCl pH 7.6, 50 mM KCl, 20 mM MgCl_2_) and then incubated for 1 h at room temperature. The remaining acetyl phosphate was removed using an ultra-filtration tube (10 KD Amicon Ultra 0.5, Millipore). 1, 3 and 10 μg phosphorylated NtrY-Kr protein were added to 200 μl of phosphorylation buffer containing 10 μg NtrXr, and incubated overnight at 28 °C. Samples were separated by a Phos-Tag^™^ Acrylamide SDS-PAGE gel (Mu Biotechnology, Guangzhou). The gel was prepared by mixing 50 μM Phostag^™^ acrylamide (29:1 acrylamide: N, N’’-methylene-bis-acrylamide) with 100 μM MnCl_2_.

Synchronized *S. meliloti* cells (Sm1021, SmLL1, Sm1021/p*ntrX* and Sm1021/p*ntrX*^D53E^) were subcultured in LB/MC broth containing 1 mM IPTG or not for 1 to 3 h. The cells from 1 ml culture were pelleted, resuspended in the buffer of 10 mM Tris-Cl, pH 7.5 and 4% SDS, incubated at room temperature for 5 min, mixed the loading dye, boiled for 10 min, and then loaded into the wells of Phos-tag^™^ acrylamide gels. Western blots were performed as described as Tang (41), with rabbit anti-NtrX (1:10000) antibodies (36). Chemiluminescent detection was performed using an ECL fluorescence colorimetric kit (Tiangen) and fluorescent signals were visualized using a Bio-Rad Gel Doc XR. Band intensities were evaluated by Image J (42).

To determine the protein levels between Sm1021 and SmLL1, synchronized cells were subcultured in 100 ml of LB/MC broth at 28 °C for half to three hours. The *ntrX* depleted cells, the synchronized cells of Sm1021/p*ntrX* and Sm1021/p*ntrX*^D53E^ were subcultured in 100 ml of LB/MC broth containing 1 mM IPTG for one to three hours. ~ 10^8^ cells were collected by centrifugation every half or one hour for each strain. 1 mg His-fused CtrA and GcrA proteins were purified through Ni^+^ columns from supernatant of *E. coli* BL21 lysates. Rabbit anti-CtrA and anti-GcrA polyclonal antibodies were prepared by Hua’an Biotech, Hangzhou.

### Microscopy

A 5-μl aliquot of fresh *S. meliloti* culture (OD_600_ = 0.15) was placed on a glass slide and covered with a cover glass. The slide was slightly baked near the edge of the flame of an alcohol lamp for a few seconds, and observed under a phase contrast microscope (Zeiss). The cells carrying pHC60 (35) were observed in GFP mode, and the images were acquired using a CCD camera Axiocam 506 color (Zeiss). The exposure time was set to 10 ms in order to capture bacterial morphology. Scanning electron microscopy was performed as described by Wang (31) to further observe cell shapes of *S. meliloti* at the mid-log phase.

### DNA sequencing and analysis

ChIP-Seq was performed by BGI (43). DNA library was prepared including DNA-end repair, 3’-dA overhang, ligation of methylated sequencing adaptor, PCR amplification and size selection (usually 100-300bp, including adaptor sequence). Bioinformatics analysis was performed as follows. The ratio of N was over 10% in whole read. Removed reads in which unknown bases are more than 10%. The ratio of base whose quality was less than 20 was over 10%. Clean Parameter: SOAP nuke filter −l 15 −q 0.5 −n 0.01 −Q 2 −c 21. After filtering, the clean data was then mapped to the reference genome by SOAP aligner/SOAP2 (Version: 2.21t). BWA (Burrows-Wheeler Aligner, Version: 0.7.10) is also used to do genome alignment after evaluating its performance. Align Parameter: soap_mm_gz −p 4 −v 2 −s 35. MACS (Model-based Analysis for ChIP -Seq, version: MACS-1.4.2): the candidate Peak region was extended to be long enough for modeling. Dynamic Possion Distribution was used to calculated p-value of the specific region based on the unique mapped reads. The region would be defined as a Peak when p-value < le-05. Peak Calling Parameter: macs14 −s 50 −g 6691694 −p 1e-5 −w --space 50 −m 10, 30. UCSC (University of California Santa Cruz) Genome Browser was used for reading peaks.

### Analysis of NtrX 3D structure

The 3D structure of *S. meliloti* NtrX was reconstructed in the server of Swiss-Model using the 4d6y template from *B. abortus* in PDB (44). The 3D structures of NtrX were analyzed by the software Pymol (Delano Scientific).

## ACKNOWLEDGMENTS

This research was supported by the National Natural Science Foundation of China (31570241 to L.L). We thanks to Dr. Yiwen Wang (Eastern Normal University) for help of SEM.

## AUTHOR CONTRIBUTIONS

L. L. designed research; S. X., L. Z., F. A., X. Y., L. H., S. Z., W. Z., and N. L. performed research; S. X., F. A., J. Y., L. Y. and L. L. analyzed data; L. L. and K. O. wrote the paper.

## REFERENCES

1. Laub MT, Shapiro L, McAdams HH. 2007. Systems biology of Caulobacter. Annu Rev Genet 41:429–41.

2. Laub MT, McAdams HH, Feldblyum T, Fraser CM, Shapiro L. 2000. Global analysis of the genetic network controlling a bacterial cell cycle. Science 290:2144–8.

3. Skerker JM, Laub MT. 2004. Cell-cycle progression and the generation of asymmetry in Caulobacter crescentus. Nat Rev Microbiol 2:325–37.

4. Panis G, Murray SR, Viollier PH. 2015. Versatility of global transcriptional regulators in alpha-Proteobacteria: from essential cell cycle control to ancillary functions. FEMS Microbiol Rev 39:120–33.

5. Jacobs C, Ausmees N, Cordwell SJ, Shapiro L, Laub MT. 2003. Functions of the CckA histidine kinase in Caulobacter cell cycle control. Mol Microbiol 47:1279–90.

6. Biondi EG, Reisinger SJ, Skerker JM, Arif M, Perchuk BS, Ryan KR, Laub MT. 2006. Regulation of the bacterial cell cycle by an integrated genetic circuit. Nature 444:899–904.

7. Jones KM, Kobayashi H, Davies BW, Taga ME, Walker GC. 2007. How rhizobial symbionts invade plants: the Sinorhizobium-Medicago model. Nat Rev Microbiol 5:619–33.

8. Van de Velde W, Zehirov G, Szatmari A, Debreczeny M, Ishihara H, Kevei Z, Farkas A, Mikulass K, Nagy A, Tiricz H, Satiat-Jeunemaître B, Alunni B, Bourge M, Kucho K, Abe M, Kereszt A, Maroti G, Uchiumi T, Kondorosi E, Mergaert P. 2010. Plant peptides govern terminal differentiation of bacteria in symbiosis. Science 327:1122–6.

9. Farkas A, Maróti G, Durgő H, Györgypál Z, Lima RM, Medzihradszky KF, Kereszt A, Mergaert P, Kondorosi É. 2014. Medicago truncatula symbiotic peptide NCR247 contributes to bacteroid differentiation through multiple mechanisms. Proc Natl Acad Sci U S A 111:5183–8.

10. Penterman J, Abo RP, De Nisco NJ, Arnold MF, Longhi R, Zanda M, Walker GC. 2014. Host plant peptides elicit a transcriptional response to control the Sinorhizobium meliloti cell cycle during symbiosis. Proc Natl Acad Sci U S A 111:3561–6.

11. Montiel J, Downie JA, Farkas A, Bihari P, Herczeg R, Bálint B, Mergaert P, Kereszt A, Kondorosi É. 2017. Morphotype of bacteroids in different legumes correlates with the number and type of symbiotic NCR peptides. Proc Natl Acad Sci U S A 114:5041–5046.

12. Barnett MJ, Hung DY, Reisenauer A, Shapiro L, Long SR. 2001. A homolog of the CtrA cell cycle regulator is present and essential in Sinorhizobium meliloti. J Bacteriol 183:3204–10.

13. Pini F, De Nisco NJ, Ferri L, Penterman J, Fioravanti A, Brilli M, Mengoni A, Bazzicalupo M, Viollier PH, Walker GC, Biondi EG. 2015. Cell Cycle Control by the Master Regulator CtrA in Sinorhizobium meliloti. PLoS Genet 11:e1005232.

14. Kobayashi H, De Nisco NJ, Chien P, Simmons LA, Walker GC. 2009. Sinorhizobium meliloti CpdR1 is critical for co-ordinating cell cycle progression and the symbiotic chronic infection. Mol Microbiol 73:586–600.

15. Pini F, Frage B, Ferri L, De Nisco NJ, Mohapatra SS, Taddei L, Fioravanti A, Dewitte F, Galardini M, Brilli M, Villeret V, Bazzicalupo M, Mengoni A, Walker GC, Becker A, Biondi EG. 2013. The DivJ, CbrA and PleC system controls DivK phosphorylation and symbiosis in Sinorhizobium meliloti. Mol Microbiol 90:54–71.

16. Xue S, Biondi EG. 2019. Coordination of symbiosis and cell cycle functions in Sinorhizobium meliloti. Biochim Biophys Acta Gene Regul Mech 1862:691–696.

17. Robinson MD, Oshlack A. 2010. A scaling normalization method for differential expression analysis of RNA-seq data. Genome Biol 11:R25.

18. Schallies KB, Sadowski C, Meng J, Chien P, Gibson KE. 2015. Sinorhizobium meliloti CtrA Stability Is Regulated in a CbrA-Dependent Manner That Is Influenced by CpdR1. J Bacteriol 197:2139–2149.

19. Pawlowski K, Klosse U, de Bruijn FJ. 1991. Characterization of a novel Azorhizobium caulinodans ORS571 two-component regulatory system, NtrY/NtrX, involved in nitrogen fixation and metabolism. Mol Gen Genet 231:124–38.

20. Nogales J, Campos R, BenAbdelkhalek H, Olivares J, Lluch C, Sanjuan J. 2002. Rhizobium tropici genes involved in free-living salt tolerance are required for the establishment of efficient nitrogen-fixing symbiosis with Phaseolus vulgaris. Mol Plant Microbe Interact 15:225–32.

21. Ishida ML, Assumpção MC, Machado HB, Benelli EM, Souza EM, Pedrosa FO. 2002. Identification and characterization of the two-component NtrY/NtrX regulatory system in Azospirillum brasilense. Braz J Med Biol Res 35:651–61.

22. Bonato P, Alves LR, Osaki JH, Rigo LU, Pedrosa FO, Souza EM, Zhang N, Schumacher J, Buck M, Wassem R, Chubatsu LS. 2016. The NtrY-NtrX two-component system is involved in controlling nitrate assimilation in Herbaspirillum seropedicae strain SmR1. Febs j 283:3919–3930.

23. Gregor J, Zeller T, Balzer A, Haberzettl K, Klug G. 2007. Bacterial regulatory networks include direct contact of response regulator proteins: interaction of RegA and NtrX in Rhodobacter capsulatus. J Mol Microbiol Biotechnol 13:126–39.

24. Lemmer KC, Alberge F, Myers KS, Dohnalkova AC, Schaub RE, Lenz JD, Imam S, Dillard JP, Noguera DR, Donohue TJ. 2020. The NtrYX Two-Component System Regulates the Bacterial Cell Envelope. mBio 11.

25. Carrica Mdel C, Fernandez I, Martí MA, Paris G, Goldbaum FA. 2012. The NtrY/X two-component system of Brucella spp. acts as a redox sensor and regulates the expression of nitrogen respiration enzymes. Mol Microbiol 85:39–50.

26. Atack JM, Srikhanta YN, Djoko KY, Welch JP, Hasri NH, Steichen CT, Vanden Hoven RN, Grimmond SM, Othman DS, Kappler U, Apicella MA, Jennings MP, Edwards JL, McEwan AG. 2013. Characterization of an ntrX mutant of Neisseria gonorrhoeae reveals a response regulator that controls expression of respiratory enzymes in oxidase-positive proteobacteria. J Bacteriol 195:2632–41.

27. Cheng Z, Lin M, Rikihisa Y. 2014. Ehrlichia chaffeensis proliferation begins with NtrY/NtrX and PutA/GlnA upregulation and CtrA degradation induced by proline and glutamine uptake. mBio 5:e02141.

28. Fernandez I, Sycz G, Goldbaum FA, Carrica MD. 2018. Acidic pH triggers the phosphorylation of the response regulator NtrX in alphaproteobacteria. Plos One 13.

29. Fernández I, Otero LH, Klinke S, Carrica MDC, Goldbaum FA. 2015. Snapshots of Conformational Changes Shed Light into the NtrX Receiver Domain Signal Transduction Mechanism. J Mol Biol 427:3258–3272.

30. Fernández I, Cornaciu I, Carrica MD, Uchikawa E, Hoffmann G, Sieira R, Márquez JA, Goldbaum FA. 2017. Three-Dimensional Structure of Full-Length NtrX, an Unusual Member of the NtrC Family of Response Regulators. J Mol Biol 429:1192–1212.

31. Wang D, Xue H, Wang Y, Yin R, Xie F, Luo L. 2013. The Sinorhizobium meliloti ntrX gene is involved in succinoglycan production, motility, and symbiotic nodulation on alfalfa. Appl Environ Microbiol 79:7150–9.

32. Calatrava-Morales N, Nogales J, Ameztoy K, van Steenbergen B, Soto MJ. 2017. The NtrY/NtrX System of Sinorhizobium meliloti GR4 Regulates Motility, EPS I Production, and Nitrogen Metabolism but Is Dispensable for Symbiotic Nitrogen Fixation. Mol Plant Microbe Interact 30:566–577.

33. De Nisco NJ, Abo RP, Wu CM, Penterman J, Walker GC. 2014. Global analysis of cell cycle gene expression of the legume symbiont Sinorhizobium meliloti. Proc Natl Acad Sci U S A 111:3217–24.

34. Khan SR, Gaines J, Roop RM, 2nd, Farrand SK. 2008. Broad-host-range expression vectors with tightly regulated promoters and their use to examine the influence of TraR and TraM expression on Ti plasmid quorum sensing. Appl Environ Microbiol 74:5053–62.

35. Cheng HP, Walker GC. 1998. Succinoglycan is required for initiation and elongation of infection threads during nodulation of alfalfa by Rhizobium meliloti. Journal of Bacteriology 180:5183–5191.

36. Zeng S, Xing S, An F, Yang X, Yan J, Yu L, Luo L. 2020. Sinorhizobium meliloti NtrX interacts with different regions of the visN promoter. Acta Biochim Biophys Sin (Shanghai) 52:910–913.

37. Schlüter JP, Reinkensmeier J, Barnett MJ, Lang C, Krol E, Giegerich R, Long SR, Becker A. 2013. Global mapping of transcription start sites and promoter motifs in the symbiotic α-proteobacterium Sinorhizobium meliloti 1021. BMC Genomics 14:156.

38. Stein BJ, Fiebig A, Crosson S. 2021. The ChvG-ChvI and NtrY-NtrX Two-Component Systems Coordinately Regulate Growth of Caulobacter crescentus. J Bacteriol 203:e0019921.

39. Szeto WW, Nixon BT, Ronson CW, Ausubel FM. 1987. Identification and characterization of the Rhizobium meliloti ntrC gene: R. meliloti has separate regulatory pathways for activation of nitrogen fixation genes in free-living and symbiotic cells. J Bacteriol 169:1423–32.

40. Van den Eede G, Deblaere R, Goethals K, Van Montagu M, Holsters M. 1992. Broad host range and promoter selection vectors for bacteria that interact with plants. Mol Plant Microbe Interact 5:228–34.

41. Tang G, Xing S, Wang S, Yu L, Li X, Staehelin C, Yang M, Luo L. 2017. Regulation of cysteine residues in LsrB proteins from Sinorhizobium meliloti under free-living and symbiotic oxidative stress. Environ Microbiol 19:5130–5145.

42. Schneider CA, Rasband WS, Eliceiri KW. 2012. NIH Image to ImageJ: 25 years of image analysis. Nat Methods 9:671–5.

43. Park PJ. 2009. ChIP-seq: advantages and challenges of a maturing technology. Nat Rev Genet 10:669–80.

44. Arnold K, Bordoli L, Kopp J, Schwede T. 2006. The SWISS-MODEL workspace: a web-based environment for protein structure homology modelling. Bioinformatics 22:195–201.

